# Selfish behavior requires top-down control of prosocial motivation

**DOI:** 10.1101/2024.01.04.574159

**Authors:** Shotaro Numano, Chris Frith, Masahiko Haruno

**Affiliations:** Center for Information and Neural Networks, NICT, 1-4 Yamadaoka, Suita, Osaka, 565-0871, Japan; Graduate School of Frontier Biosciences, Osaka University 1-3 Yamadaoka, Suita, Osaka 565-0871 Japan; Institute of Neurology, University College London, 17-19 Queen Square, London WC1N 3AZ, UK

## Abstract

Individuals must regularly choose between prosocial and proself behaviors. While past neuroscience research has revealed the neural foundations for prosocial behaviors, many studies have oversimplified proself behaviors, viewing them merely as a reward-maximization process. However, recent behavioral evidence suggests that response times for proself behaviors are often slower than those for prosocial behaviors, suggesting a more complex interdependence between prosocial and proself neural computations. To address this issue, we conducted an fMRI experiment with the ultimatum game, where participants were requested to accept (money distributed as proposed) or reject (both sides receive none) offers of money distribution. In the decisions, the participants could maximize self-interest by accepting the offer (i.e., proself), while by rejecting it, they could punish unfair proposers and promote the “equity” social norm (i.e., prosocial). We constructed a drift diffusion model (DDM) that considers both behavioral choices and response times and used the DDM parameters in our fMRI analysis. We observed that participants who suppressed inequity-driven rejection behaviors displayed heightened dACC activity in response to disadvantageous inequity. Importantly, our functional connectivity analysis demonstrated that the dACC exhibited negative functional connectivity with the amygdala when unfair offers were presented. Furthermore, the PPI connectivity encoded the average reaction time for accepting unfair offers (i.e., proself behaviors). Considering that the amygdala also responded to disadvantageous inequity in these experiments and previous studies, these results show that the top-down control of prosocial motives (i.e., aversion to disadvantageous inequity) plays a key role in implementing proself behaviors.

## Introduction

Choosing a prosocial or proself behavior constitutes a fundamental aspect of human social hehavior^1,2^. Research has identified the brain mechanism for prosocial behaviors exerting cognitive control of the selfish drive^3–6^. For instance, several studies reported that low frequency repetitive (suppressive) transcranial magnetic stimulation to the right dorsolateral prefrontal cortex^3–5,7^ (dlPFC) decreases human prosocial behavior, suggesting that the dlPFC suppresses self-interest to achieve prosocial behaviors. Other studies have identified prosocial motives represented in subcortical structures. It was reported that the ventral striatum and amygdala^8–14^, together with the dorsal anterior cingulate cortex (dACC), encode prosocial motives such as the aversion to iniquity. Recent animal studies also reported that the amygdala and ACC (or prelimbic region in rodents)^15–17^ are crucial for cooperative behaviors in non-human primates and rodents.

In contrast with prosocial behavior, researchers have proposed a simple reward-maximization process as the neural underpinning of proself behaviors^2,18,19^. More recently, however, Rand and colleagues found that prosocial behaviors are often faster than proself behaviors^20^. The response time difference in prosocial and proself behaviors becomes larger when an individual is under time pressure^20–22^ or making decisions under a cognitive load^23–25^.

Slower response times for proself behaviors compared with prosocial behaviors^20^ may imply a more complex neural computation is at work for implementing proself behaviors. We hypothesized that top-down control of intuitive prosocial motives underlies the selection of proself behaviors. To test this hypothesis, we conducted functional magnetic resonance imaging (fMRI) of the ultimatum game^11,26,27^ (**Fig. 1**) and used a drift-diffusion model (DDM) (**Fig. 1c**) to analyze behavioral choices and reaction times simultaneously.

**Fig. 1.**
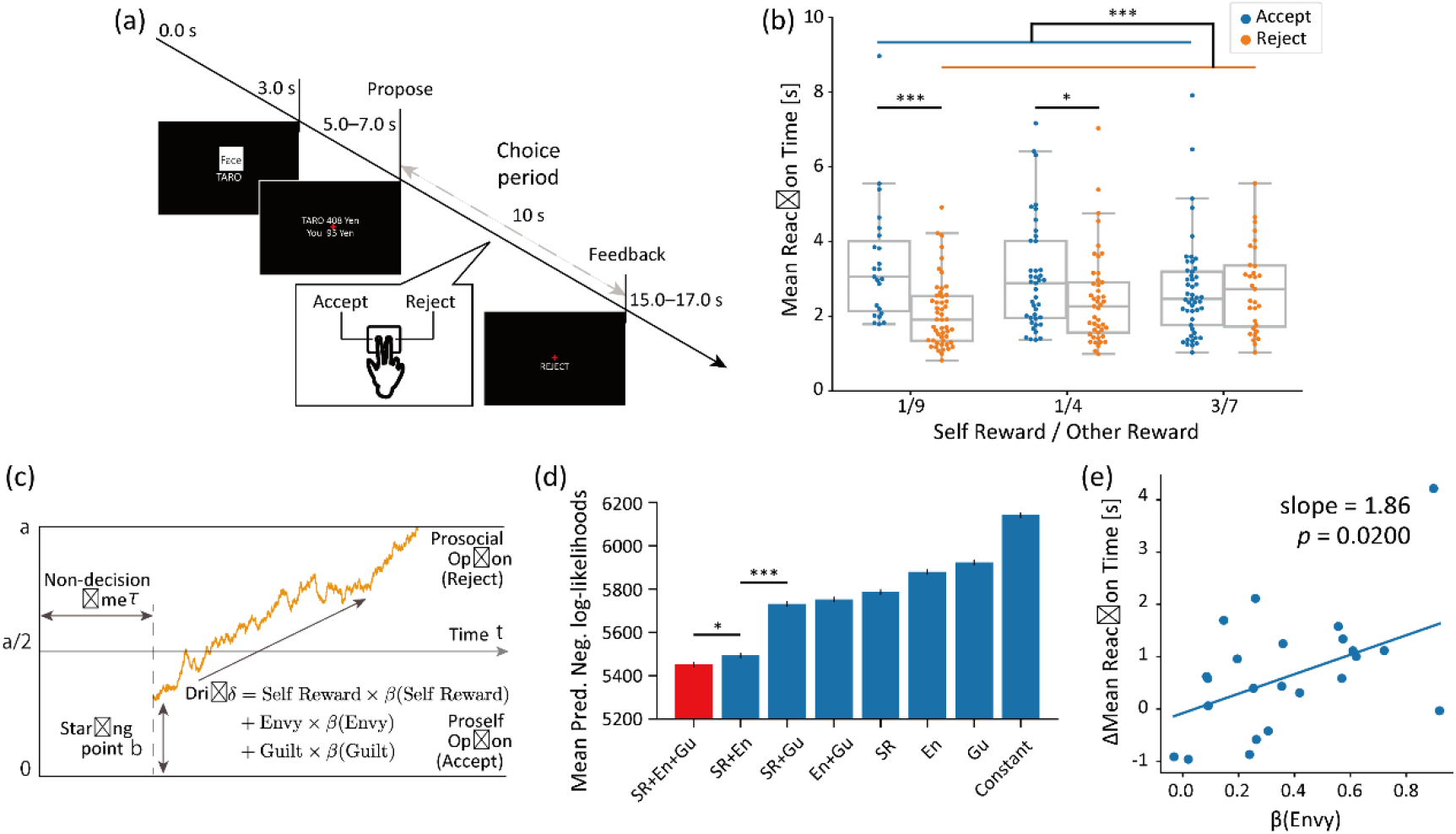
The task and computational model. **(a)** The ultimatum game. In each trial, a participant was shown a proposer’s face (not shown in this version) and name, and presentation of a money distribution offer followed. The participant was required to accept or reject the offer and report his/her choice by pressing a button within 10 s. If the participant accepted the offer, the money was distributed as proposed, while neither received any amount if the offer was rejected. **(b)** Mean reaction times of each participant for disadvantageous offers are separately displayed for accept (blue) and reject (orange) choices (*t*(72) = 4.50 and \*\*\**p* = 2.57 × 10^−5^, *t*(84) = −2.02 and \**p* = 0.0466, *t*(238) = 3.57 and \*\*\**p* = 4.31 × 10^−4^, Student’s t-test, respectively). **(c)** The drift-diffusion model (DDM). The orange trajectory represents a participant’s decision processes in a trial. The participant makes a decision (i.e., prosocial (reject) or proself (accept) choice to an offer) when the trajectory reaches either the upper or lower boundary. Our full DDM model includes four components: a distance between the upper and lower boundaries *a*; a relative starting point *b*; a drift rate *δ*, represented as a linear combination of the self reward (SR), disadvantageous inequity (DI) and advantageous iniquity (AI); and a non-decision time *τ*. **(d)** The model selection. Predictive negative log-likelihoods for possible models. Each bar represents the mean negative predictive log-likelihoods (*t*(3996) = 2.42 and \**p* = 0.0155, *t*(3997) = −13.6 and ∗∗∗ *p* = 2.15 × 10^−41^, Welch’s *t*-test, respectively). Red shows the best model SR+DI+AI, in which the *δ* term is defined as SR × *β*(SR) + DI × *β*(DI) + AI × *β*(AI). In the other models, *δ* is a combination of no more than two of SR, DI, and AI. “Constant” represents the model whose *δ* is a constant. Eu, envy; Gu, guilt. **(e)** Correlation between (DI) and individual differences between the mean reaction times for acceptance and for rejection. Each point represents a participant who tended to accept a disadvantageous offer (robust linear regression with Tukey’s bi-weight function; slope = 1.86 and *p* = 0.0200).

The DDM is a computational model for binary decision-making^28–30^ that assumes a decision maker accumulates information sequentially and executes a decision as soon as the decision process reaches either of the two boundaries. Moreover, a DDM can estimate the model parameters that best explain both the behavioral choice and response time data, as shown in many human^31–35^ and animal^35–38^ studies utilizing a wide range of tasks from perceptual decisions^39–45^ to social behavior^22,30,31,46–51^.

## Results

### Effects of disadvantageous inequity on choices and reaction times

Healthy participants (n=58) underwent an fMRI scan while playing an ultimatum game^26^ as a responder (**Fig. 1a**). In this game, proposers made a series of offers about money sharing to the responder, who decided whether to accept or reject each offer within 10 seconds. If the responder accepted an offer, the money was distributed as proposed; otherwise, neither side received any money. Each proposal consisted of pairs of rewards for the participant (self-reward; SR) and the proposers (other-reward; OR).

The rejection of unfair offers in the ultimatum game can be seen as a form of altruistic punishment designed to increase equity by punishing the behavior of the unfair proposer^52^. Altruistic punishment is an example of prosocial behavior, because the costly rejection by the responder may contribute to the benefit of the group if the proposer behaves more fairly in the future. By contrast, the acceptance of unfair offers can be seen as a form of proself behavior because it maximizes the participant’s self-interest.

Human behavior in the ultimatum-game is reported to be influenced by disadvantageous inequity (DI) and advantageous inequity (AI) separately (both defined using SR and OR; see Computational models for inequity-aversion in the Methods). We therefore first confirmed that participant behaviors were consistent with prior findings regarding DI and then examined the relationship between the choice and response time.

The mean rejection rate for a disadvantageous offer (i.e., SR/OR < 1/1) was 46.1% (s.d. 31.5%; **Sup. Fig. 1**), while that for an advantageous offer (i.e., SR/OR ≧ 1/1) was 2.81% (s.d. 7.08%). The difference between the two conditions was significant (*t*(58) = 12.9, *p* = 2.13 × 10^−20^, paired *t*-test). A repeated one-way analysis of variance (ANOVA) of the mean rejection rates for disadvantageous offers yielded a significant main effect of the SR/OR rate (*F*(3, 316) = 1.01 × 10^2^, *p* < 8.19 × 10^−41^). The mean rejection rates in the strongest (i.e., SR/OR = 1/9) and weakest (i.e., SR/OR = 2/3) DI trials were 79.2% (s.d. 35.3%) and 17.0% (s.d. 30.2%), with the former significantly higher than the latter (*t*(58) = 13.8 and *p* = 6.37 × 10^−22^, paired *t*-test). These results clarified that participants rejected more as the degree of DI increased.

We next examined the participants’ decision times (response time) (**Fig. 1b** and **Sup. Fig. 2**). The mean reaction time to disadvantageous offers was 2.341 s (s.d. 0.8631 s) and to advantageous offers it was 1.765 s (s.d. 0.6758 s). The mean reaction time for acceptance was significantly longer than the one for rejection in both the 1/9 and 1/4 offers (see Ultimatum Game in the Methods for an explanation about the offers; **Fig. 1b**; *t*(72) = 4.50 and \*\*\**p* = 2.57 × 10^−5^, *t*(84) = −2.02 and \**p* = 0.0466, Student’s *t*-test, respectively). We also found a significant difference between the reaction times for acceptance and rejection (*t*(238) = 3.57 and \*\*\**p* = 4.31 × 10^−4^, Student’s *t*-test) of disadvantageous offers. By contrast, for advantageous offers, the mean reaction time was significantly longer for the rejection than for the acceptance (3/2 and 7/3; **Sup. Fig. 2**; *t*(58) = −2.80 and *** *p* = 0.0702, *t*(63) = −6.12 and *** *p* = 6.68 × 10^−8^, Student’s *t*-test). These results confirmed that only for disadvantageous offers were acceptances (proself choices) slower than rejections (prosocial choices).

To explore the neural mechanism for slower proself behaviors, we constructed DDM models (**Fig. 1c** and **Methods**) and selected the best one to predict behavior (**Fig. 1d**). More specifically, we calculated the predictive negative log-likelihood for eight possible models using the repeated stratified eight-fold cross-validation. This computation process was repeated 2000 times, and we calculated the mean predictive negative log-likelihood (NLL) for each model. The best model with the lowest NLL included three components in the drift term: SR+DI+AI (predictive NLL = 1.09 × 10^7^). However, the model using SR+DI was not statistically different from the best model (*t*(3996) = 2.42 and *p* = 0.155, Welch’s *t*-test), and these two models had a much lower NLL than the other models (*t*(3997) = −13.6 and *p* = 2.15 × 10^−41^, Welch’s *t*-test), showing that DI and SR in the drift term are important for predicting the choice and response time.

We conducted a correlation analysis between individual parameter of the best model and the mean rejection rate and the mean reaction time of the same individual. *β*(DI), which represents the DDM coefficient for DI in the drift term, showed the strongest correlation with both the mean rejection rate (Pearson’s *r* = 0.689 and *p* = 3.22 × 10^−16^) and the mean reaction time (Pearson’s *r* = −0.4099 and *p* = 1.40 × 10^−3^). Further, we found that *β*(DI) represents individual differences between the mean reaction times for acceptance and rejection (**Fig 1e**; robust linear regression with Tukey’s bi-weight function; slope = 1.86 and *p* = 0.0200) of disadvantageous offers.

### Neural activity correlated with Disadvantageous Inequity

Having established that *β*(DI) accurately represents behavioral choices and response times in unfair conditions, we used it in our fMRI analysis. First, we conducted a within-participant (first level) general linear model (GLM) analysis for the disadvantageous-offer onsets and performed a group-level (second level) one-sample *t*-test to replicate the previous studies for DI^27^. We observed strong activations in the bilateral anterior insula and dACC (**Sup. Fig. 3** and **Sup. Table 1**; *p* < 0.05 cluster-level family-wise error (FWE) corrected; the cluster-formation height threshold was *p* < 0.001). These results showed not only that the dACC and anterior insula are important in processing DI, but also that our experiment successfully captured brain activity patterns reported in previous studies^11,27^.

We then conduced a parametric fMRI analysis based on the DDM parameters. We hypothesized for disadvantageous offers, accepting behavior is realized by top-down control (or suppression) of the aversion to DI. We therefore sought the brain structure whose activity is correlated with DI at the individual level and beta values of the group correlated with −*β*(DI). The assumption behind this analysis was that the top-down control process should be stronger when DI is large and even stronger in participants who accept more disadvantageous offers (i.e., correlated with - *β*(DI)). We identified activity in the dACC and posterior cingulate cortex (PCC) (**Fig. 2a** and **Table 1**; *p* < 0.05 cluster-level FWE corrected). This result suggested that the cingulate area play a crucial role in top-down control of DI aversion.

**Fig. 2.**
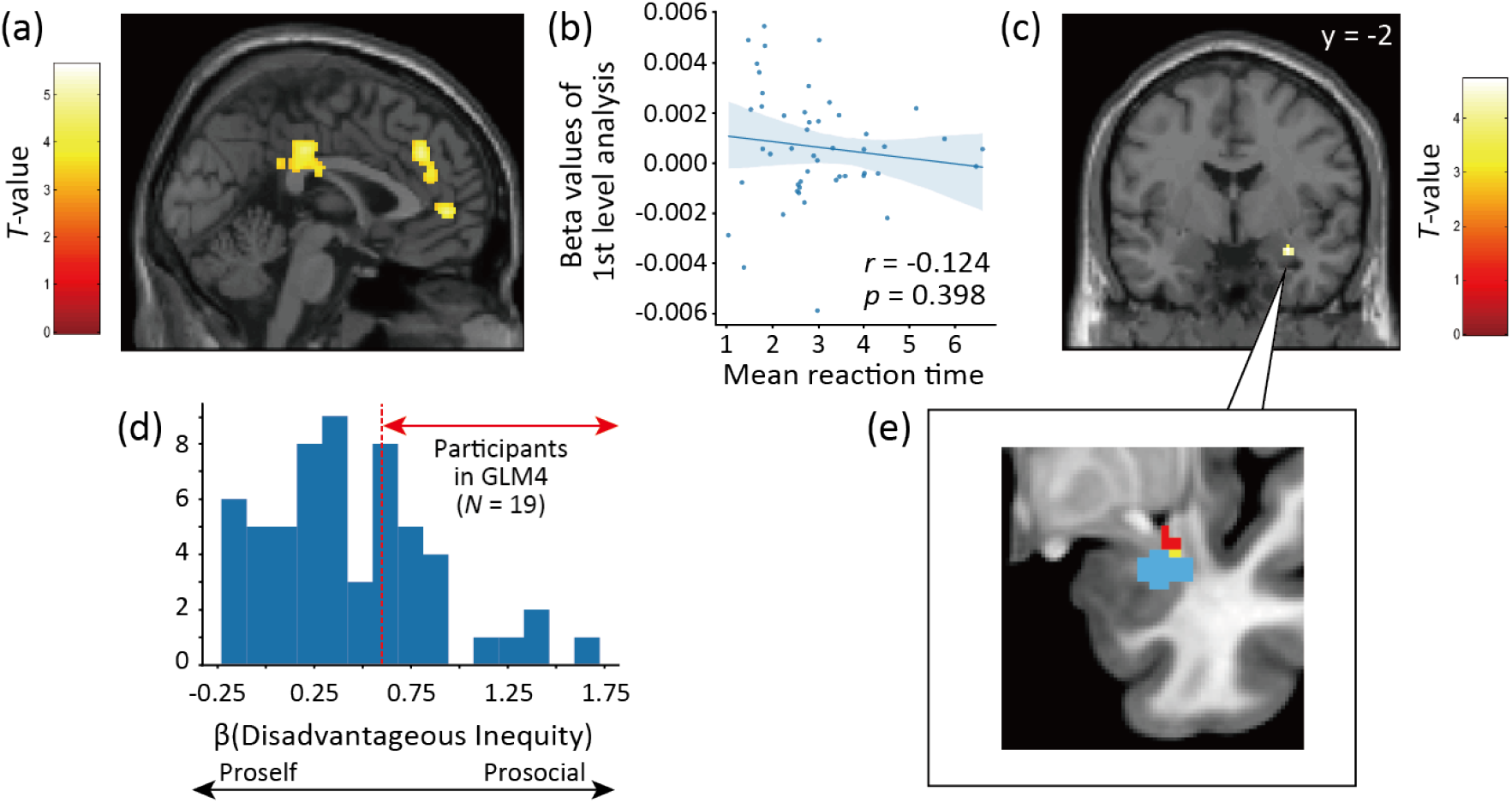
Negative functional connectivity between the dACC and the amygdala upon Disadvantageous Inequity in proself participants. **(a)** dACC activity correlated with DI for proself participants (MNI coordinates [4, 34, 32], GLM2, *p* < 0.05, cluster-level FWE corrected). The first-level analysis used DI as a parametric modulator, and the second-level analysis used −*β*(DI) as the covariate. Because a large and small *β*(DI) represents the prosocial and proself tendency of a participant, respectively, the activity suggests that the dACC responds to disadvantageous offers more in proself participants. **(b)** Correlation between individual beta values in the dACC and individual mean reaction times for the acceptance of disadvantageous offers was not significant (Pearson’s *r* = −0.124, *p* = 0.398). **(c)** A PPI analysis of dACC activity (see **a**) and DI (experimental factor) identified negative connectivity with the amygdala (MNI coordinates [30, −2, −18]). This finding suggests that when DI is large, dACC activity suppresses the amygdala. **(d)** The distribution of *β*(DI). **(e)** Overlap in the amygdala between the PPI results (**Fig.2c** blue) and amygdala activity in response to DI (red) in prosocial participants (red, N=19, shown in **d**). The yellow voxels represent the overlap. (*p* < 0.05, small volume corrected for the amygdala ROI.)

**Table 1.** Neural activities in response to Disadvantageous Inequity weighted by. −(DI). *P* < 0.05 cluster-level FWE corrected.

| Brain area | MNI coordinates | | | Voxel size (k) | $t$ value |
| --- | --- | --- | --- | --- | --- |
| | $x$ | $y$ | $z$ | | |
| Occipital Inf L | -20 | -96 | -6 | 489 | 5.63 |
| Occipital Inf R | 28 | -92 | -4 | 470 | 5.58 |
| Middle Frontal Gyrus R | 24 | 62 | 26 | 201 | 5.47 |
| Dorsal Anterior Cingulate Cortex R | 4 | 34 | 32 | 861 | 5.31 |
|  | -12 | 34 | 34 |  | 4.85 |
| Dorsal Anterior Cingulate Cortex | 0 | 50 | 2 | 260 | 4.75 |
| Posterior Cingulate Cortex | 2 | -26 | 36 | 360 | 5.14 |
| SupraMarginal R | 54 | -46 | 42 | 538 | 5.09 |
| Cerebellum Crus2 L | -36 | -66 | -40 | 463 | 4.79 |
| Supp Motor Area R | 14 | 22 | 64 | 264 | 4.57 |
| Parietal Inf L | -56 | -48 | 36 | 235 | 4.19 |

Next, we speculated that the correlation between dACC activity and *β*(DI) may account for the slower reaction times for accepting disadvantageous offers. Accordingly, we conducted a correlation analysis between individual beta values in the dACC and the mean reaction times. However, we did not find a significant correlation (**Fig. 2b**), suggesting that the dACC is not directly responsible for slower prosocial behavior.

We also investigated neural activity with AI in addition to DI. Although we applied the first-level results of AI regression to a second-level analysis contrasting *β*(AI), we found no significant correlation with *β*(AI) (*p* < 0.05 cluster-level FWE corrected), potentially reflecting a smaller number of AI trials.

### dACC-amygdala interaction encodes response times for accepting disadvantageous offers

To further examine the top-down control mechanism from the cingulate areas displayed in **Fig. 2a**, we conducted a psychophysiological interaction (PPI) analysis using the gPPI toolkit^53^. We defined a volume of interest from the peak positions of each cingulate cluster and ran PPI by setting these regions as the seed. We looked for negative interactions.

We found a significant negative interaction between the dACC (MNI coordinates [−12, 34, 34]; uncorrected *p* < 0.001; cluster size k>10) and the amygdala (MNI coordinates [30, −2, −18], **Fig. 2c** and **Table 2**). Notably, we did not use any DDM-derived parameters to reveal this connectivity because DDM parameters may produce false-positive correlations with response times. We also performed the same PPI analysis for other cingulate peaks (MNI coordinates [4, 34, 32] and [0, 50, 2] in the dACC, and [2, - 26, 36] in the PCC; see also **Table 1**) but did not find any significant negative correlation at uncorrected *p* < 0.001. These results suggest that the dACC activity correlated with DI is involved in the top-down control of amygdala activity.

**Table 2.** Neural activity identified by the PPI analysis. Uncorrected *P* < 0.001 and cluster extent threshold *k* > 20.

| Brain area | MNI coordinates | | | Voxel size (k) | $t$ value |
| --- | --- | --- | --- | --- | --- |
| | $x$ | $y$ | $z$ | | |
| Amygdala R | 30 | -2 | -18 | 28 | 4.71 |

Next, we examined whether amygdala activity also correlates with DI in prosocial participants. We tested the participants who behaved in a prosocial manner (largest number of rejections), i.e., largest *β*(DI) (*N* = 19; **Fig. 2d**). Indeed, we found a cluster activity in the amygdala to DI that was consistent with previous studies^8–14^. This cluster partially overlapped with the amygdala cluster in **Fig. 2e** and **Sup. Fig. 4** (*p* < 0.05 small-volume FWE corrected).

We then investigated if this negative interaction between the dACC and amygdala encodes the response times for accepting disadvantageous offers. We found that the dACC-amygdala connectivity exhibits a strong correlation with the response time for accepting disadvantageous offers (**Fig. 3a**; Pearson’s *r* = 0.349 and *p* = 0.00905) but not advantageous offers (**Fig. 3b**; Pearson’s *r* = 0.163, *p* = 0.222). These results suggest that the dACC underlies the acceptance of DI by cognitive control of the DI aversion encoded in the amygdala. In other words, through this interactive process, prosocial urges can be suppressed.

**Fig. 3.**
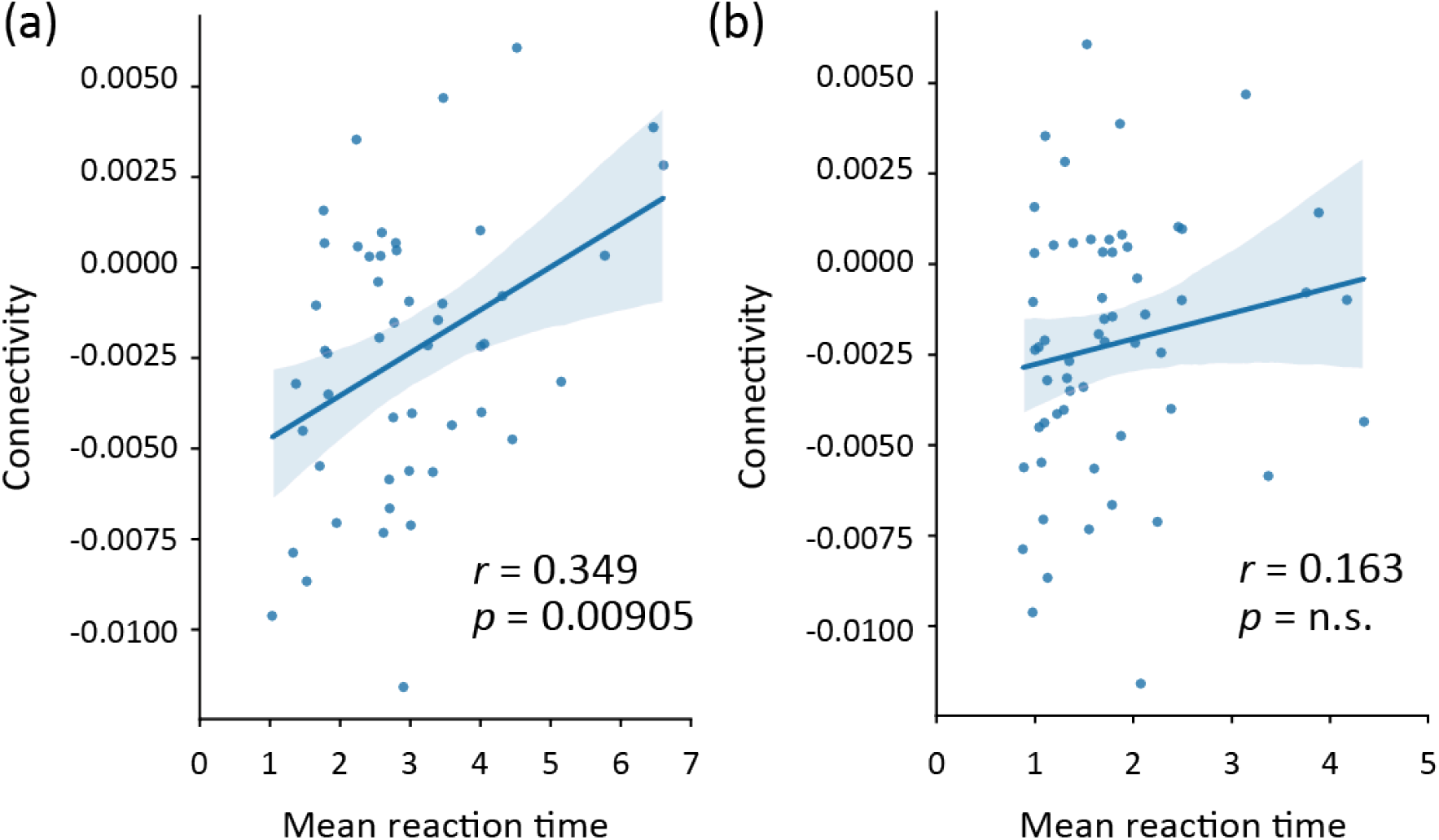
PPI connectivity between the dACC and amygdala predicts mean reaction times for proself behavior. **(a)** The correlation between individual PPI connectivity and individual mean reaction times for accepting disadvantageous offers was significant (Pearson’s *r* = 0.349, *p* = 0.00905). **(b)** By contrast, there was no such correlation for advantageous offers (Pearson’s *r* = 0.163, *p* = 0.222).

### Comparing DDM and standard value-based model

Finally, we compared the DDM and a standard value-based model (i.e., a logistic model) that considers choices but not reaction times. We analyzed the participants’ behavior and neural activity by the logistic model. For this purpose, we assumed that the best predictive logistic model has the same three parametric components as the best DDM model.

We estimated the coefficients *γ* of the logistic model using R^54^ and found a significant correlation between *γ*(DI) and the mean rejection rate (Pearson’s *r* = 0.740 and *p* = 3.28 × 10^−11^). However, the correlation between *γ*(DI) and mean reaction time was not significant (Pearson’s *r* = −0.206 and *p* = 0.120). Furthermore, *γ*(DI) did not show a positive correlation with the difference in mean reaction times for acceptance and rejection (**Fig 4a**; robust linear regression with Tukey’s bi-weight function; slope = 0.467 and *p* = 0.0951), which is in contrast to *β*(DI) (**Fig. 1e**). These data confirmed that only the DDM can capture behaviors in the time domain.

**Fig. 4.**
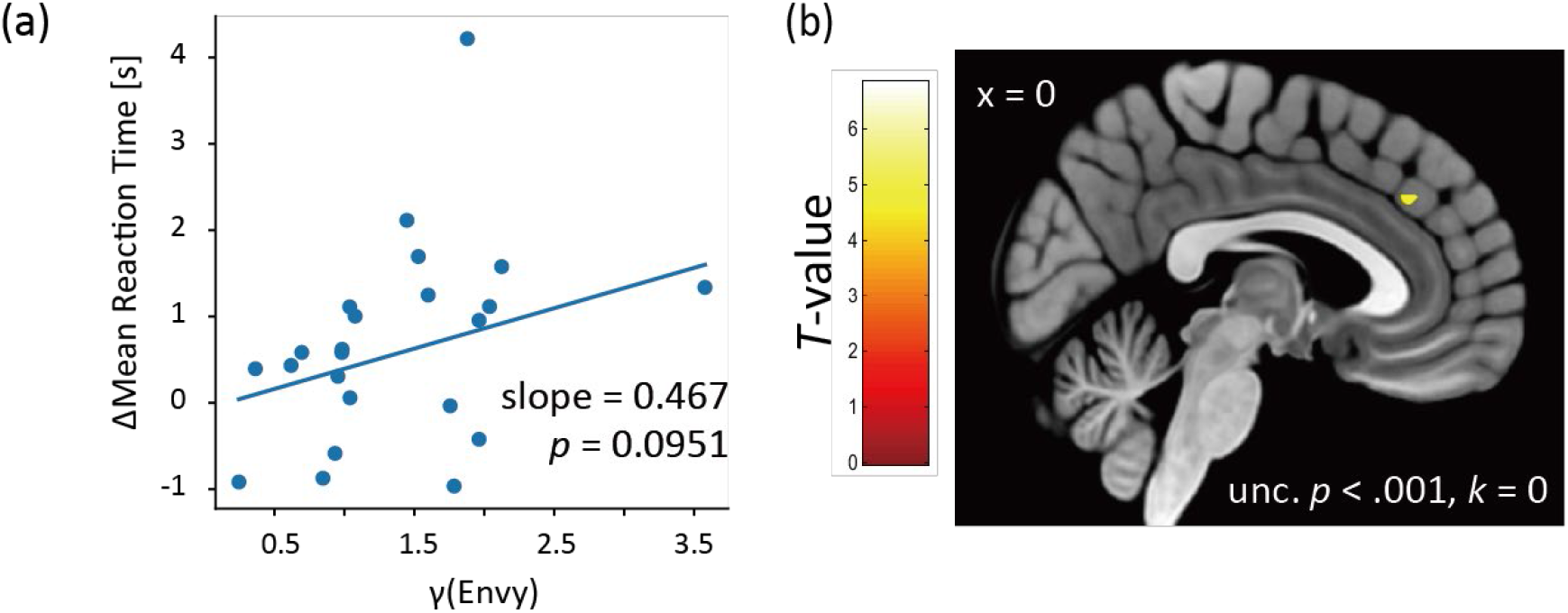
Comparisons of our DDM and a value-based decision model. **(a)** Linear regression between *γ*(DI) and individual differences between mean reaction times for acceptance and rejection. Each point represents an acceptive participant (robust linear regression with Tukey’s bi-weight function; slope = 0.467 and *p* = 0.0951). **(b)** A second-level analysis was conducted with γ(DI), the weight for DI in the value-based model (uncorrected *p* < 0.001). Only weak activity in the dACC activity was observed even with a moderate statistical threshold.

Observing that only *β*(DI), but not *γ*(DI), encodes response times **(Fig. 1e** and **Fig. 4a**), we compared the two parameters in the fMRI analysis. More concretely, we performed a GLM analysis, where all second-level regressors were replaced by the coefficients of the logistic model. In sharp contrast with **Fig. 2a**, we identified much weaker activity in the dACC (**Fig. 4b** and **Sup. Table 2**; *p* < 0.001 uncorrected). These results suggest that the dACC is involved in computing the choice and response time and that the DDM is more suitable for detecting dACC functions.

## Discussion

This study investigated why proself behavior is often slower than prosocial behavior. To investigate this question, we conducted fMRI on participants performing the ultimatum game and analyzed the behavioral and fMRI data with a DDM that considers behavioral choices and response times. We found that top-down control of prosocial motivations plays a key role in implementing proself behavior and accounts for the slower response time. More specifically, we found that participants who exhibited a smaller DDM parameter for DI (i.e. *β*(DI)) and dampened disadvantage-driven rejection (i.e., higher acceptance) showed heightened dACC activity for DI. Our PPI analysis revealed that, when unfair offers are presented, the dACC exhibits a negative functional connectivity with the amygdala, which also responds to DI. We demonstrated that PPI connectivity between the dACC and amygdala encodes the average reaction time for accepting unfair offers but not other offers. We also demonstrated that a logistic regression model that only considers choices performed poorly compared with the DDM, indicating that the consideration of both the choice and response time benefits elucidation of the neural mechanism for proself behavior.

The amygdala^8–14^ and dACC are known to be involved in the aversion to social equity and conflict monitoring^55,56^. As shown in **Figure 2b**, dACC activity did not correlate with the response time for accepting disadvantageous offers. Instead, PPI connectivity between the dACC and amygdala encoded the response time. This link between functional connectivity and response time indicates that the dACC suppresses the amygdala response to disadvantageous inequity after detecting the accept/reject conflict. This view is consistent with previous anatomical and physiological studies on animals, which found strong projections from the dACC to the amygdala, particularly to its basolateral part (BLA), in both humans^57^ and non-human primates^58–60^. Additionally, a recent study of rodents engaged in social interactions reported that prelimbic (regarded as an analogue of human dACC) projections to the BLA contributes to self-interest motives for decision-making^17^.

Previous studies reported that top-down control of selfish drives is a key mechanism for implementing prosocial behavior^3–6^. The present study demonstrated that top-down control of the amygdala response by the dACC mechanisms is also important for proself behavior, possibly changing the traditional view of proself and prosocial behaviors.

A natural question arises regarding whether the dACC suppresses the amygdala response to inequity directly through a single projection or indirectly through an internal circuit within the amygdala, since the amygdala mainly receives inputs from the BLA and outputs the processed information from the central nucleus of the amygdala and other nuclei^58,61^. Although it is difficult to address this question with the spatial resolution of fMRI, we think the latter is more likely considering the limited overlap between PPI activity and inequity activity in the amygdala (**Fig. 2e**). In addition, we identified functional connectivity from the left dACC to the right amygdala. A previous report suggested that the amygdala has different functions in the left and right hemispheres^62^. However, further studies are needed to determine what this asymmetry represents.

The present study may also provide clues on the link between prosocial/proself behaviors and psychiatric disorders. For instance, research has suggested that individuals with psychopathy may have a dysfunctional amygdala, which could compromise their prosocial behaviors^63^. The results from the present study potentially indicate different selection mechanisms for the proself behavior between healthy and psychopath populations. That is, due to this amygdala dysfunction, psychopaths may have never acquired an aversion to inequity, which underlies prosocial motivation. In addition, the dACC-amygdala interaction we report in this study may be absent in the proself behaviors of autistic and depressed population^12^.

In this study, the source of top-down control was within an individual`’`s brain. However, arguments for controlling prosocial motivations often arise from the cultural context. For example, we may be easily persuaded that “greed is good”. The top-down control mechanism by dACC reported here likely plays a central role in addressing how such ideas are instantiated in bran function.

This study utilized the ultimatum game. The rejection of unfair offers in this game can be seen as a form of altruistic punishment designed to increase equity by punishing the behavior of the unfair proposer^52^. Altruistic punishment is an example of prosocial behavior because costly rejection by the responder can contribute to the benefit of the group if the proposer behaves more fairly in the future. On the other hand, the acceptance of unfair offers can be seen as a form of proself behavior because it maximizes the participant’s self-interest. By using DI in our DDM, we focused on proself aspects in the ultimatum game. However, recent behavioral studies^64,65^ comparing multiple economic game tasks suggested that rejections in the ultimatum game do not correlate with cooperation in other games and caution may be necessary when using the number of rejections as a general measure of prosocial preferences. Therefore, a series of functional neuroimaging experiments using other economic games are recommended to determine the range of proselfness to which our findings can be generalized. Nonetheless, the present study updates our understanding of human prosocial/proself behaviors by showing that top-down control of prosocial motives plays a key role in implementing proself behavior, accounting for its longer response time.

## Methods

### Participants

We used G*power^66^ (version 3.1.9.3) to calculate the optimal sample size by setting the following parameters: “Test family” as “T test”; “Statistical test” as “Linear multiple regression: Fixed model, *R*^2^ deviation from zero”; “Type of power analysis” as “A priori”; “Effect size *f*^2^” as 0.2; “*⍺* err prob” as 0.05; “Power (1 − *β* err prob)” as 0.9; and “Number of predictors” as 3. We obtained the output parameters “Total sample size” as 55 and “Actual power” as 0.902, based on which we determined the number of participants in the main experiment.

In the main experiment, we recruited 68 participants and excluded 10 due to their large and/or jaggy head movements (more than 1 mm) in their fMRI data, leaving 58 participants (36 males and 22 females; mean age = 22.41, s.d. = 1.702) for the main experiment. The protocol of this study was approved by the National Institute of Information and Communications Technology’s ethics committee. Written informed consent was obtained from all participants.

### Experiment procedure

Participants first read the task procedure and filled a consent form. They were then told “The proposals in the task were made by students at a university nearby, and the proposer and the proposal are different from trial to trial. Your responses will affect both the proposer’s and your rewards”. In addition, the participants were instructed to decide within ten seconds and press a corresponding button as soon as possible. Before starting the task, the participants experienced the time limit (< 10s) and task setting with three practice trials. After the main task, we interviewed all participants. They also completed a social value orientation survey, which we did not use in the present study.

### Ultimatum game

Participants performed the ultimatum game (**Fig. 1a**) as a responder. They evaluated the proposals from an anonymous proposer and decided to accept or reject the proposal by pressing a button within 10 seconds. Error trials with late responses (longer than 10 s) were excluded from the analysis. If a responder accepted a proposal, money was distributed as proposed. If not, no money was received by either the proposer or responder.

Base offers were one of: ¥350–150, ¥300–200, ¥250–250, ¥200–300, ¥150–350, ¥100–400 or ¥50–450 for the participant and proposer. The displayed offers were constructed by adding a uniform random number ranging from ¥-25 to ¥25 in each trial. This design was to encourage participants to think and avoid pattern-based responses. There were eight trials for each based offer; therefore, 56 trials in total.

In each trial, after three seconds of fixation, a proposer’s face and name were presented for one second. Participants then had 10 seconds to accept or reject a proposal (choice period in **Fig. 1a**), and their choice was shown for 1 second (“Feedback” phase in **Fig. 1a**). After the feedback, the participant rested for 6.5 seconds before the next trial. Each trial took 22.5-24.5 seconds, and the entire task took 22 minutes and 16 seconds. In the main experiment, the viewing distance was 91.0-94.0 cm, the viewing angle was 21.7-22.4 degrees, and the screen luminance was 110.3 cd/m^2^.

### Computational models for inequity-aversion

We defined disadvantageous inequity (DI) and advantageous inequity (AI) at trial *t* as:

> DI_*t*_ = max(other-reward_*t*_ − self-reward_*t*_, 0)

> AI_*t*_ = max(self-reward_*t*_ − other-reward_*t*_, 0),

where Self Reward (SR)_*t*_ is the reward for the responder (participant) and Other Reward (OR)_*t*_ is the reward for the proposer. To focus on proself behavior, we investigated how DI, AI and SR influence a participant’s rejection of unfair offers and their response time by using two computational models: a drift-diffusion model (DDM) and a logistic model.

The DDM is characterized by four parameters: a decision boundary, relative starting point, non-decision time, and drift rate δ (**Fig. 1c**). The decision boundary indicates the amount of information needed to decide, which varies depending on the difficulty of the task and how deliberate the participant makes decisions. In other words, it represents the level of caution. An increase in the boundary results in fewer errors but slower responses. The relative starting point represents the response bias of each participant. If the participant prefers option A, they quickly decide for A since the amount of information required for A is lower than for option B. The non-decision time represents the time for peripheral processes, such as encoding the stimulus and moving a hand to press a key. Finally, δ indicates whether the participant’s decision is toward option A or B as time goes by. A larger δ yields a quicker and more robust decision. Consequently, the total reaction time is the sum of the time to diffuse from the starting point to the upper or lower boundary and the non-decision time.

In this study, we associated the prosocial option (i.e., rejection) with the upper bound of the DDM and the proself option (i.e., acceptance) with the lower bound. We describe δ as:

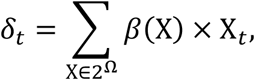

where *β*(X) is the weight of variable X, and 2^Ω^ is the power set of variables in the task: Ω = {SR, AI, DI}. To complete the parameter estimation process within a reasonable time, we set the evidence height (*a*) and non-decision time (*τ*) to be constant across all participants; this approach is used in the EZ method^67,68^.

We conducted a preliminary experiment using the same task, recruiting 13 participants (seven males and six females; mean age = 20.8, s.d. = 2.01) and determined *a and τ*. We estimated these two parameters for the optimal model and used the medians as the parameter values in the main experiment (*a* = 4.240, *τ* = 0.549). We implemented the maximum likelihood estimation method for the other two parameters (the relative starting point and δ) using the Nelder-Mead method by utilizing the likelihood function provided by the RWiener package^69^.

Next, we defined a subjective value of prosocial choice _{*t*,prosocial}_ and a subjective value of proself choice *V*_{*t*,proself}_ at trial *t* for the logistic model. We integrated these values into a value (difference) function (*t*) for a participant as follows:

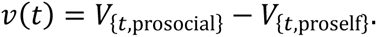

This value function provides the probability of choosing the prosocial option *p*_prosocial_(*t*) by using a logistic function: *p*_prosocial_(*t*) = 1⁄[1 + exp(−*v*(*t*))]. To compare the DDM and the value-based logistic model, we assumed that the best predictive logistic model has the same components as the drift term in the best predictive DDM. We therefore re-defined the subjective value of prosocial and proself choice as follows:

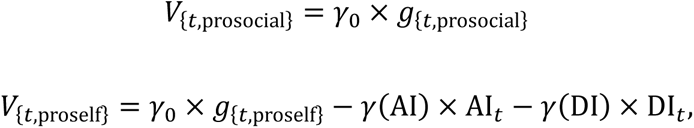

where *g*_{*t*,X}_ is a participant’s gain for option X, and *γ* is a coefficient. _{*t*,prosocial}_ was 0 and *g*_{*t*,proself}_is SR*t* in our task. We conducted the logistic regression analysis using the bias-reduction method provided by the Brglm package^70^.

### Model selection

Since we defined the drift rate δ as a linear combination of variables in the task, we have eight possible models:

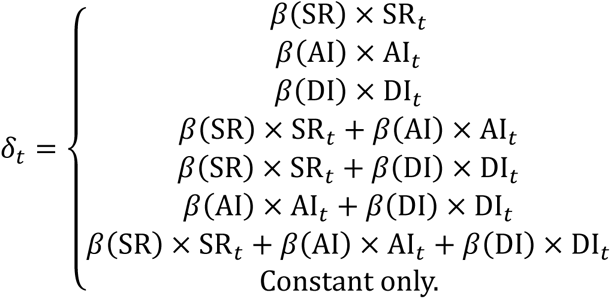

We identified the best predictive model based on the repeated *k*-fold method. We randomly divided all trials into seven folds to ensure that each fold includes seven SR/OR conditions and estimated each *β* using seven out of the eight folds for a given model. We then calculated the predictive log-likelihood for the remaining fold using the estimated *β*. We repeated this procedure 2000 times, computed the sum of the predictive negative log-likelihoods, and chose the best model with the smallest sum.

### fMRI acquisition

We examined participants’ neural activity during the task performance by fMRI. Neural images were acquired using a 3T MRI (Prisma; Siemens Medical Systems) with a 64-channel head coil at the Center for Information and Neural Networks, National Institute of Information and Communications Technology. To correct for distortion, we first acquired field maps covering the whole brain with the following parameters: repetition time [TR] = 753 ms, echo time 1 [TE1] = 5.16 ms, echo time 2 [TE2] = 7.62 ms, flip angle [FA] = 9°, field of view [FOV] = 256 mm, and voxel size [VS] = 2×2×2 mm. We then acquired T2*-weighted images during a task using a multi-echo planner pulse sequence with the following parameters: TR = 2000 ms, TE = 30 ms, FA = 75°, FOV = 200 mm, VS = 2×2×2 mm, and multi-band factor = 3. We finally acquired high-resolution T1-weighted images after the task session using the MPRAGE pulse sequence with the following parameters: TR = 1900 ms, TE = 3.37 ms, FA = 9°, FOV = 256 mm, and VS = 1×1×1 mm.

### Image preprocessing

Images were processed using SPM12 (http://www.fil.ion.ucl.ac.uk/spm) on MATLAB 2018a. Image preprocessing included slice-timing correction (reference time, 1000 ms), motion correction, removal of movement-induced variance (unwarping) with a measured field map, co-registration to T1 image, spatial normalization, and spatial smoothing with a 6 mm full width at half maximum (FWHM). We conducted these procedures with the default parameters of SPM12 except the reference time in slice-timing correction and FWHM size in spatial smoothing. After preprocessing, we found that we needed to exclude 10 participants who had large (> 1 mm) and/or jaggy head movements.

### General linear model analysis

We conducted individual-level and group-level general linear model (GLM) analyses. In the individual-level analysis, we constructed a design matrix that consists of 14 regressors. The first regressor represented the offer event onset timing. The second to fourth regressors were parametric modulators: DI, SR and AI. The fifth regressor was the proposer’s name and face presentation onset timing. The sixth and seventh regressors were button-press and feedback onset timing. The eighth to thirteenth regressors represented the participant’s head movements, and the fourteenth regressor represented a constant. The event duration of the first to fourth regressors was set to a participant’s reaction time in each trial, and the fifth to the seventh regressors was set to 0 s. We performed the GLM analysis using these design matrices with the following default parameter: high pass filter cutoff = 128 s, and the basis function was set as a canonical hemodynamic response function.

In the group-level analysis, we used contrast images associated with a parametric modulator of DI in the participant-level analysis. We constructed a factorial design to investigate neural activity correlated with DI. We investigated the results with cluster-level family-wise error (FWE) corrected *p* < 0.05 (cluster-forming height threshold was *p* < 0.001).

We also constructed two multiple regression designs to investigate neural activity correlated with the parameters of the DDM. The first design included the participant’s *β*(DI) and *β*(SR) and the second one included *γ*(DI) and *γ*(SR). We included an intercept in each design, and each covariate was not standardized. We investigated the results with cluster-level FWE corrected *p* < 0.05 (cluster-forming height threshold was *p* < 0.001).

### Functional connectivity analysis

To reveal the neural mechanism for DI, we investigated functional connectivity from the seed peaks in **Fig. 2a** using a psycho-physiological interaction (PPI) analysis. We extracted the BOLD signals from the participant-level GLM results to conduct the PPI analysis. Focusing on the anterior-dorsal (da) ACC area, we set the statistical height threshold at *p* < 0.05 corrected for multiple comparisons at the cluster level (the cluster-forming height threshold of *p* < 0.001) and found two peaks in the daACC area. We defined the volume of interest (VOI) as a sphere that had a 4 mm radius centered at these daACC peak positions (MNI coordinates were [−12, 34, 34] and [4, 34, 32]).

We conducted the PPI analyses using the generalized PPI toolbox^53^ implemented on SPM12. We first constructed two participant-level PPI designs whose seed-ROIs were at [−12, 34, 34] and [4, 34, 32] based on the design matrix used in the GLM analysis. We performed the participant-level PPI analyses and then conducted one-sample *t*-tests with positive and negative contrasts in the group-level PPI analysis.

### Functional connectivity and reaction time

We were interested in the relationship between proself choices (i.e., accepting unfair offers) and their reaction times. Any functional connectivity involved in this link is likely correlated with the reaction times and rejection rate. To examine this hypothesis, we extracted the peak intensity (beta value) of the PPI connectivity whose cluster size was larger than 10 (*i.e*., a voxel-size over 80 mm^3^) and conducted a correlation analysis between the individual peak intensity of the PPI connectivity and the individual average reaction times for accepting unfair offers.

