## Supplementary figures and tables for "Selfish behavior requires top-down control of prosocial motivation"

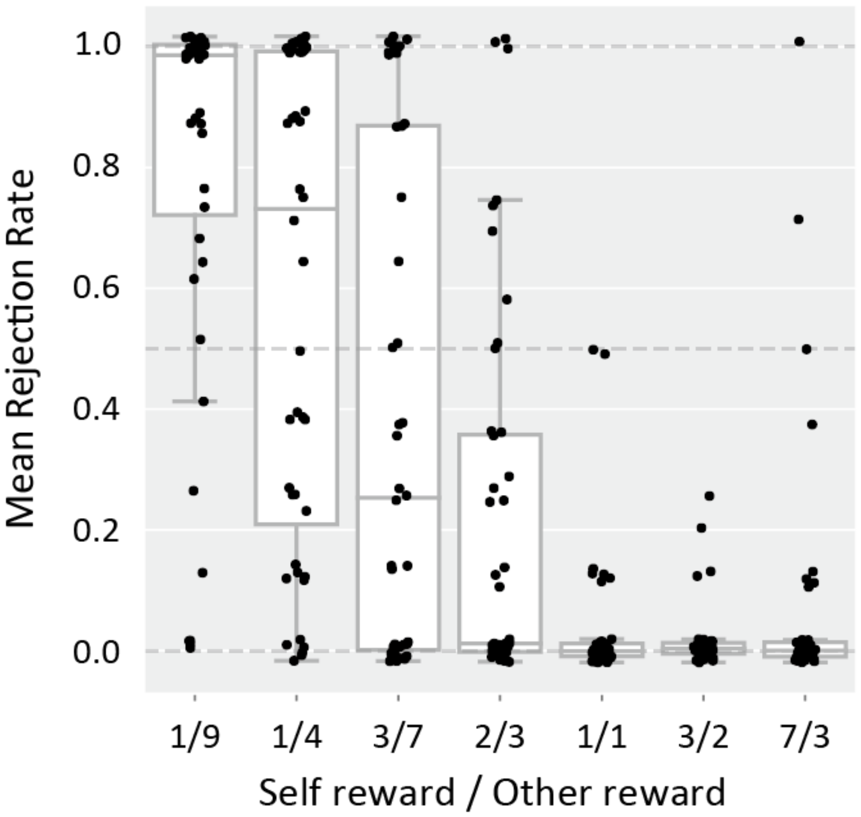


**Supplementary Figure 1. Mean rejection rates for different offers.**

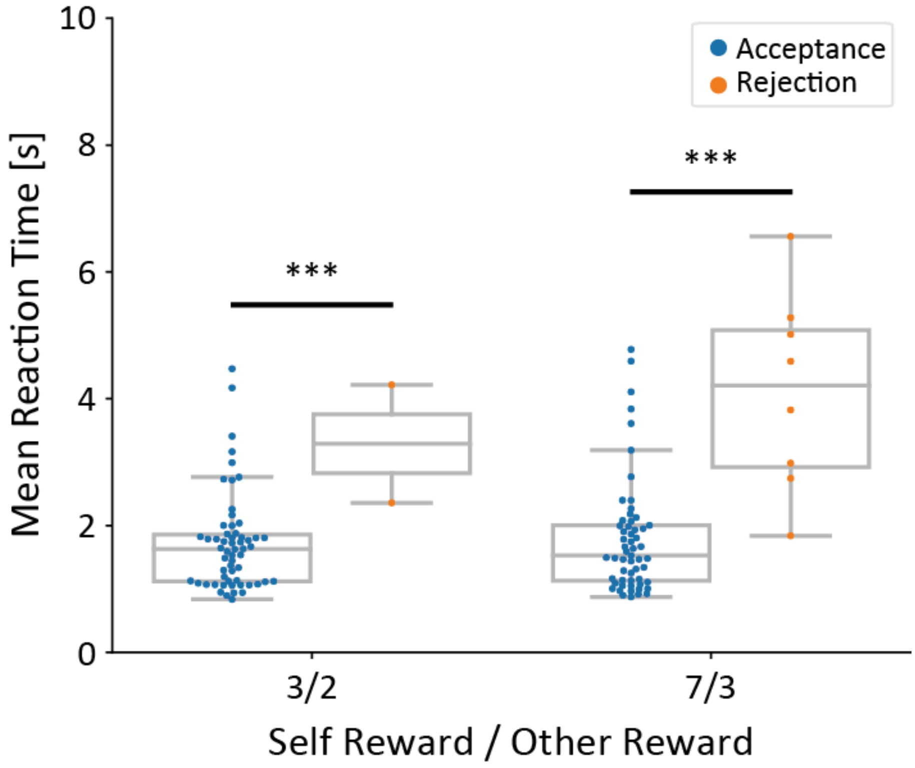


**Supplementary Figure 2. Mean reaction times for advantageous offers.** Blue and orange dots represent the mean reaction times for accept and reject, respectively ($t(58) = -2.80$ and $p = 0.00702$, and $t(63) = -6.12$ and $p = 6.68 \times 10^{-8}$, Student's *t*-test, respectively).

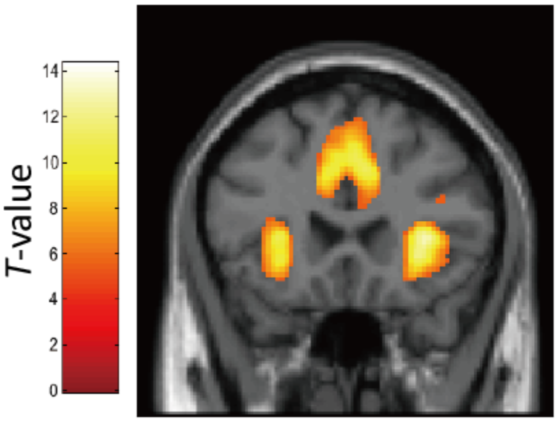


**Supplemental Figure 3. Neural activations in Disadvantageous Inequity conditions** (p<0.05, peak-level family-wise error (FWE) corrected).

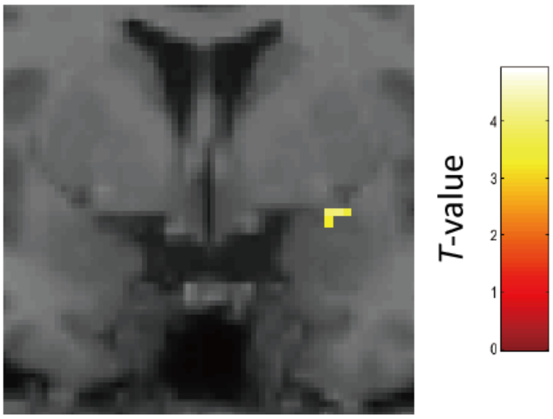


**Supplemental Figure 4. Amygdala activity correlated with Disadvantageous Inequity in prosocial participants** ($p < 0.005$, small-volume corrected for the amygdala ROI).

**Supplementary Table 1. Neural activities for Disadvantageous Inequity conditions.** $P < 0.05$ cluster-level FWE corrected; cluster extent threshold $k > 10$.

| Brain area | MNI coordinates *x* | *y* | *z* | Voxel size (k) | *t* value |
|---|---|---|---|---|---|
| Insula R | 34 | 20 | 8 | 13040 | 14.3 |
| Thalamus L | -6 | -20 | 0 | 2924 | 11.4 |
| Dorsal Premotor Cortex L | -48 | 8 | 30 | 589 | 10.1 |
| Angular R | 26 | -58 | 46 | 2282 | 9.91 |
| Vermis 9 | 0 | -58 | -34 | 1604 | 9.84 |
| Hippocampus R | 24 | -36 | -2 | 213 | 8.90 |
| Cerebellum 6 L | -30 | -56 | -32 | 905 | 8.58 |
| Cerebellum 8 R | 14 | -62 | -50 | 180 | 7.89 |
| Inferior Frontal Gyrus R | 42 | 30 | 20 | 156 | 7.46 |
| Pallidum R | 12 | 2 | -4 | 278 | 7.30 |
| Calcarine R | 12 | -68 | 10 | 130 | 6.88 |
| Calcarine L | -14 | -74 | 8 | 128 | 6.80 |
| Middle Frontal Gyrus L | -28 | 44 | 30 | 33 | 5.86 |
| Middle Frontal Gyrus R | 28 | 42 | 28 | 35 | 5.78 |

**Supplementary Table 2. Neural activities correlated with Disadvantageous Inequity in the value-based model ($\boldsymbol{\gamma}(\mathbf{DI})$).** $P < 0.05$ cluster-level FWE corrected.

| Brain area | MNI coordinates *x* | *y* | *z* | Voxel size (k) | *t* value |
|---|---|---|---|---|---|
| Occipital Mid L | -26 | -98 | 0 | 812 | 6.79 |
| Occipital Inf R | 26 | -96 | -4 | 498 | 5.78 |
| Parietal Inf R | 50 | -58 | 48 | 394 | 4.96 |
| Cerebellum Crus2 L | -34 | -66 | -40 | 370 | 4.67 |
| Superior Frontal Gyrus R | 10 | 30 | 60 | 223 | 4.38 |
